# AllerStack: Predicting Allergenic Proteins with a Stacked Ensemble Approach

**DOI:** 10.1101/2025.06.22.660976

**Authors:** Harshika Sharma, Mansi Goel, Devkul Sahu, Mazhar Ayaz Sayed, Pratham Mangla, Pratham Shekhawat, Shubham Yadav, Ganesh Bagler

## Abstract

Accurate prediction of protein allergenicity is essential for ensuring food and drug safety. While machine learning and deep learning models have been explored for this task, limitations remain in dataset scale, feature representation, and model architecture. Here, we introduce AllerStack, a two-stage stacked ensemble model integrating handcrafted and ESM2-based learned features for allergenicity classification. The model was developed using a balanced dataset comprising 11,930 allergenic and 11,930 non-allergenic proteins. We extracted amino acid composition, dipeptide composition, and physicochemical features using the Biopython library, alongside contextual embeddings from the pre-trained ESM2 protein language model. Diverse classifiers (QDA, SVM, KNN, and ANN) were trained separately on these features in the base layer. Their predictions were used as input to a meta-classifier based on XGBoost. AllerStack achieved high predictive performance with 96.87% accuracy, 96.86% F1-score, 93.75% Matthews correlation coefficient (MCC), and an AUC of 0.99. A publicly accessible web server (https://cosylab.iiitd.edu.in/allerstack/) enables real-time allergenicity prediction from protein sequences. AllerStack provides a robust, interpretable, and user-friendly platform for allergen detection in computational biology.

**Highlights:**

- An extensive dataset of 23,860 proteins.
- Combines handcrafted and ESM2-derived features.
- Stacked ensemble model with an accuracy of 96.87%.
- SHAP-based model interpretability at the model and feature level.
- Public web server, AllerStack (https://cosylab.iiitd.edu.in/allerstack/).

## 1. Introduction

Allergic diseases are a growing global health concern, affecting nearly 20% of the population and contributing to chronic conditions such as asthma, rhinitis, dermatitis, and respiratory distress [1,2]. These diseases result from aberrant immune responses to allergens, such as chemical substances, aeroallergens, or food proteins. While proteins are vital for nutrition and cellular function, specific proteins, such as those found in peanuts, shellfish, and gluten, can trigger severe allergic reactions in some individuals, leading to issues like anaphylaxis, respiratory distress, or gastrointestinal problems. Computational biology offers a powerful approach to addressing these challenges by enabling the prediction of allergenic proteins through data-driven modeling. By integrating machine learning with molecular biology and nutritional science, researchers can analyze large-scale protein datasets to identify patterns associated with allergenicity. These predictive models help uncover the underlying determinants of immune reactivity and support the identification of safer protein sources for food and therapeutic applications.

The underlying immune mechanism involves the production of immunoglobulin E (IgE) antibodies, which sensitize mast cells to release inflammatory mediators like histamine and cytokines upon re-exposure [3]. The increasing prevalence and healthcare burden of allergic diseases underscore the urgent need for robust allergen screening tools [4]. Proteins with at least six consecutive matching amino acids or more than 35% sequence similarity to known allergens are considered potential allergens, according to the Food and Agriculture Organization. However, these threshold-based approaches often overlook structural and physicochemical determinants of allergenicity. Computational models have emerged to address these limitations by leveraging machine learning (ML) and deep learning (DL) for pattern recognition in biological sequences.

Several methods have been proposed for allergenicity prediction. AlgPred 2.0 integrates BLAST and MERCI motif search, achieving an AUC of 0.98 [5]. With a similar spirit, AllerHunter employs SVM-based pairwise comparisons across 1,356 allergens to identify cross-reactivity [6]. AllerCatPro extends this by incorporating 3D epitope mapping with 84% accuracy [7]. Alignment-free models have also been developed to overcome limitations of sequence similarity. AllerTop applies the k-nearest neighbors (KNN) algorithm using ACC-transformed features from 2,210 allergens [8]. AllergenFP, another alignment-free tool, leverages a descriptor-based fingerprint approach to identify allergens and non-allergens [9].

Recent DL approaches have further enhanced predictive performance. ProAll-D employs LSTM on a 2,427-sequence dataset with an AUC of 91.5% [10]. In a similar spirit, BERT-based model by Wang et al. achieved an AUC of 95% using transformer and ensemble techniques on food allergens [11]. Other notable contributions include DeepAlgPro (accuracy: 91.24%) [12], ALLERDET (97.26%) [13], and hybrid frameworks like AllerHybrid, which combine ANN and SVM models using k-mer and physicochemical features, attaining 94.28% accuracy [14]. Researchers implemented a deep learning-based ensemble method that uses Extra Tree, Deep Belief Network, and CatBoost models to identify protein allergens [15].

Despite these advancements, challenges remain in scaling models to larger datasets, integrating interpretable features, and generalizing across diverse protein families. We present AllerStack, a stacked ensemble framework that fuses classical feature engineering and deep contextual protein embeddings to address these gaps. Our model is trained on a balanced and curated dataset of 11,930 allergenic and 11,930 non-allergenic proteins. Features include amino acid composition, dipeptide composition, physicochemical descriptors (via Biopython), and embeddings from the transformer-based ESM2 model. It integrates predictions with multiple base classifiers (QDA, SVM, KNN, ANN) and an XGBoost meta-classifier. The ensemble approach captures complex sequence-to-function relationships across multiple feature domains. Our best-performing configuration achieves an accuracy of 96.87% and an AUC of 0.99. Furthermore, a publicly accessible web server (https://cosylab.iiitd.edu.in/allerstack/) supports end-user prediction from raw sequences.

## 2. Materials and Methods

### 2.1. Dataset Collection

A comprehensive dataset of allergenic proteins was constructed by integrating multiple publicly available and validated sources. Allergenic protein sequences were obtained from COMPARE (2,784) [16], AllergenOnline (2,291) [17], ADFS (2,204) [18], IUIS (1,109) [19], ALLERMATCH (391) [20], ALLERBASE (1,737) [21], SDAP (1,887) [22], and Swiss-Prot (1,288), using the query “allergen AND reviewed: yes” [23].

### 2.2. Data Preprocessing

Sequences lacking UniProt identifiers or without literature validation were removed. Entries containing non-natural amino acids (B, O, J, U, X, Z) were excluded. After refinement, 11,930 allergenic sequences were retained, incorporating filtered entries from the databases above.

Since proteins explicitly labeled as “non-allergen” are not available, we randomly selected 11,930 non-allergenic sequences from Swiss-Prot using the query “NOT allergenic AND reviewed: yes” [24], resulting in a balanced dataset of 23,860 sequences.

### 2.3. Redundancy Removal and Dataset Partitioning

To minimize redundancy and avoid bias due to sequence similarity, clustering was performed using CD-HIT with a 90% sequence identity threshold [25,26]. Allergenic and non-allergenic sequences resulted in 2,158 and 4,829 clusters, respectively. These clusters were split into training and external validation sets using an 80:20 split. The training data is further divided into training and internal validation subsets. Stratified sampling ensured equal representation of allergenic and non-allergenic sequences in each split.

### 2.4. Feature Extraction

Feature extraction transforms protein sequences into discrete data of a defined length. Three main types of features were extracted: composition-based, physicochemical-based, and deep contextual embeddings.

#### 2.4.1 Composition-Based Features

Amino acid composition (AAC) and dipeptide composition (DPC) were computed using Biopython v2.11 [27]. AAC generates a 20-dimensional vector representing residue frequency, while DPC produces a 400-dimensional vector by quantifying dipeptide frequencies.

#### 2.4.2 Physicochemical-based Features

Physicochemical descriptors were derived using Pseudo Amino Acid Composition (PAAC), considering composition and sequence [28]. Additionally, molecular weight, aromaticity, isoelectric point (pI), instability index, and secondary structure fractions (alpha-helices, beta-sheets, coils) were calculated via Biopython [27]. These attributes relate to protein stability, solubility, and immunogenic potential [29–31].

#### 2.4.3 ESM2

We extracted 1,280-dimensional embeddings using Meta AI’s pre-trained ESM (Evolutionary Scale Modeling 2) model [32]. The contextualized representations were obtained by averaging transformer-layer outputs across amino acid tokens, capturing high-level biological information [33].

### 2.5. Feature Scaling and Selection

All features were scaled to prevent variables with large magnitudes from dominating model performance. GradientBoosting-based feature selection was used to reduce dimensionality and select the most informative features [34,35]. This approach reduced the risk of overfitting and enhanced the model’s generalization ability.

### 2.6. Stacked Ensemble Model Framework

We initially evaluated multiple machine learning models on Biopython features, including Quadratic Discriminant Analysis (QDA), Linear Discriminant Analysis (LDA), Extreme Gradient Boosting (XGBoost), Light Gradient Boosting Machine (LGBM), Adaptive Boosting (AdaBoost), Random Forest (RF), RidgeClassifierCV (RCV), and Extremely Randomized Tree (ExtraTree) with five-fold cross-validation. ESM2-derived features were evaluated using Extremely Randomized Tree (ExtraTree), Support Vector Classifier (SVC), K-nearest neighbors (KNN), RCV, LGBM, and deep learning models: Artificial Neural Network (ANN), Recurrent Neural Network (RNN), and Long Short Term Memory (LSTM). The most effective models were selected to form the final stacked ensemble pipeline.

The final stacked model consists of four base classifiers: QDA, ANN, KNN, and SVC, followed by an XGBoost meta-classifier trained on their predictions [36–38]. The architecture of the stacked model is depicted in Figure 1.

**Figure 1:**
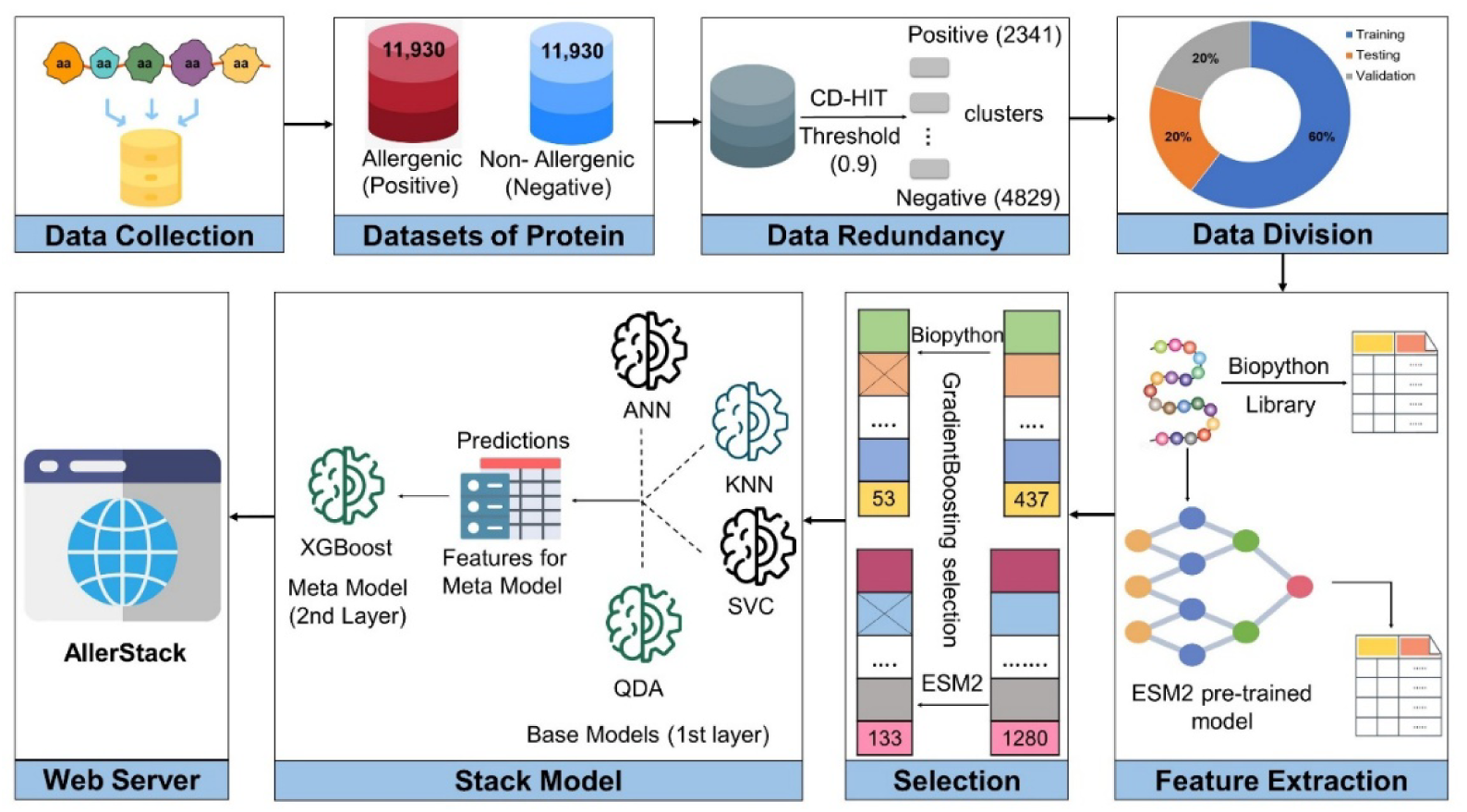
Schematic representation of the AllerStack framework. The architecture includes four base classifiers (QDA, ANN, KNN, and SVC), whose outputs are used by a meta-classifier (XGBoost) for final allergenicity prediction.

### 2.7. Evaluation Metrics

Model performance was assessed using standard classification metrics: Sensitivity (SENS), Specificity (SPEC), Precision (PREC), Accuracy (ACC), Matthews Correlation Coefficient (MCC), and F1 score, and Area under the Receiver Operating Characteristic curve (AUC-ROC) used in many previous studies [5,10,39,40]. These metrics capture both threshold-dependent and threshold-independent performances, enabling robust evaluation of binary classification under balanced and imbalanced conditions.

## 3. Results

### 3.1. Compositional Analysis

We analyzed the amino acid composition of allergenic and non-allergenic proteins to identify residue-level trends associated with allergenicity. Our study showed that amino acids such as Aspartic Acid (D), Glutamic Acid (E), Isoleucine (I), Lysine (K), Leucine (L), and Glutamine (Q) were more prevalent in allergenic sequences, suggesting a potential role in eliciting immune responses. The enrichment of negatively charged residues like D and E may facilitate immunogenic interactions, thereby enhancing allergenic potential [41]. Sicherer and Sampson also reported that hydrophobic amino acids such as I and L can contribute to increased protein stability and persistence within the host, potentially prolonging immune system exposure[42].

Conversely, non-allergenic sequences exhibited higher frequencies of Cysteine (C), Phenylalanine (F), Glycine (G), Asparagine (N), Arginine (R), Serine (S), Threonine (T), and Tryptophan (W). Prior studies have shown that residues like C and N are commonly involved in glycosylation and disulfide bond formation, which influence protein folding and may reduce immunogenicity [43]. Additionally, amino acids such as R and T have been implicated in maintaining protein solubility and structural integrity, which may further reduce allergenic potential [44]. Supplementary Figure 1 illustrates the average amino acid composition differences between allergenic and non-allergenic proteins.

### 3.2. Physicochemical Analysis

We evaluated a range of physicochemical properties of allergenic and non-allergenic proteins, including molecular weight, sequence length, aromaticity, isoelectric point (pI), instability index, secondary structure composition (alpha-helices, beta-sheets), acidity, basicity, hydrophobicity, hydrophilicity, flexibility, net charge at pH 7, and sequence complexity, as shown in Supplementary Figure 2.

Allergenic proteins generally exhibit higher molecular weight and longer sequences, characteristics that may contribute to increased structural complexity and allergenic potential [45]. These proteins also display greater aromaticity, indicating enrichment of aromatic residues such as phenylalanine, tyrosine, and tryptophan, which are known to stabilize protein structures through hydrophobic interactions and π-π stacking [46,47].

The pI values of allergenic proteins are typically higher, potentially influencing their interactions with immune receptors and antibody binding [48]. Allergenic proteins also tend to have lower instability indices, suggesting enhanced structural stability and resistance to degradation [49]. In terms of secondary structure, allergenic proteins exhibit balanced distributions of alpha-helices and beta-sheets, which may contribute to overall structural integrity [50]. The distribution of acidic and basic residues influences solubility and immune recognition, and allergenic proteins show greater variability. Similarly, their hydrophobic character is elevated, which can promote aggregation and enhance immunogenic persistence [42,51].

Increased flexibility in allergenic proteins may facilitate conformational changes that expose epitopes to immune cells. Furthermore, a higher net charge at physiological pH and elevated sequence complexity may enhance immunogenic potential by increasing epitope diversity and immune engagement [44,52].

### 3.3. Motif Analysis

To explore conserved sequence patterns characteristic of allergenic and non-allergenic proteins, we performed motif discovery using STREME from the MEME Suite [53]. Motif analysis helps reveal biologically meaningful residue patterns that may be associated with allergenicity and immune recognition. These motifs, shown in Figure 2, highlight the discriminative sequence patterns between allergenic and non-allergenic proteins.

**Figure 2:**
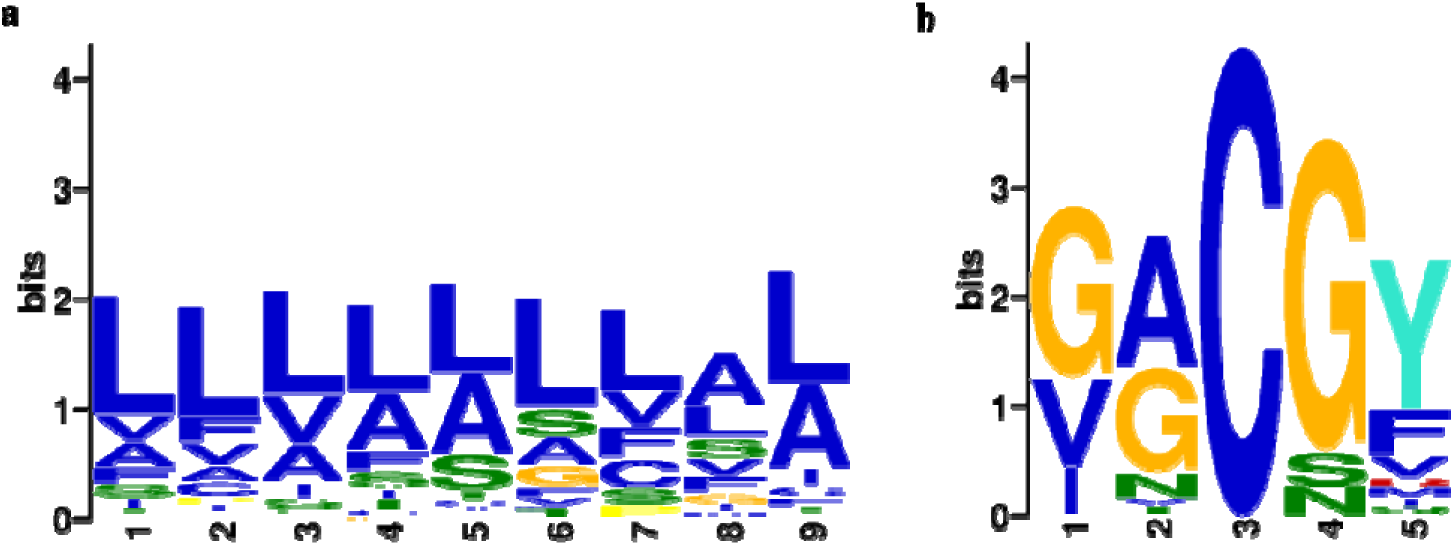
Allergenic and non-allergenic motif logos identified using STREME from the MEME suite. (a) Allergens: motif LLLLLLLAL (p = 6.3 × 10^-5^) identified in 30.5%. (b) Non-allergens: motif GACGY (5.8 × 10^-4^) found in 25.5%.

In allergenic proteins, STREME identified a highly conserved 9-mer motif: LLLLLLLAL, present in 30.5% of sequences, with a statistically significant p-value of 6.3 × 10^-5^. This motif is dominated by hydrophobic residues, particularly Leucine (L) and Alanine (A), suggesting a possible role in maintaining protein stability or forming epitope-rich regions that facilitate immune recognition. The prevalence of Leucine-rich segments has been previously linked to structural rigidity and resistance to proteolysis traits commonly associated with allergenic proteins.

In contrast, non-allergenic proteins were enriched for a distinct 5-mer motif: GACGY, occurring in 25.5% of sequences with a p-value of 5.8 × 10^-4^. This motif includes a mix of polar (Glycine, Cysteine) and acidic (Aspartic acid) residues, which may contribute to flexible and hydrophilic regions that are less likely to engage IgE-mediated immune responses. The presence of Cysteine (C) in this motif could also indicate a role in disulfide bond formation, potentially enhancing structural stability without eliciting allergenic responses.

### 3.4. Performance of Traditional Machine Learning Models

We evaluated nine traditional machine learning (ML) classifiers: Quadratic Discriminant Analysis (QDA), Linear Discriminant Analysis (LDA), XGBoost, Light Gradient Boosting Machine (LGBM), AdaBoost, Random Forest (RF), RidgeClassifierCV (RCV), Extremely Randomized Trees (ExtraTree), and Ridge Regression to distinguish between allergenic and non-allergenic sequences. Each model was optimized through hyperparameter tuning, and their performance wa assessed on both test and validation datasets, as shown in Table 1. Among these, QDA achieved the highest classification performance, with an accuracy of 90.83% on the test set and 91.26% on the validation set. Its F1 scores were 89.78% and 88.23%, respectively, indicating reliable prediction capabilities.

**Table 1:** Performance of traditional machine learning classifiers on Biopython-derived features across test and validation datasets. Metrics include sensitivity (SENS), specificity (SPEC), precision (PREC), accuracy (ACC), Matthews correlation coefficient (MCC), and F1 score.

| Model | Test Data |  |  |  |  |  | Validation Data |  |  |  |  |  |
| --- | --- | --- | --- | --- | --- | --- | --- | --- | --- | --- | --- | --- |
|  | SENS | SPEC | PREC | ACC | MCC | F1 | SENS | SPEC | PREC | ACC | MCC | F1 |
| QDA | 95.66 | 92.07 | 93.22 | 90.83 | 84.75 | 89.78 | 94.52 | 91.23 | 94.23 | **91.26** | 86.58 | 88.23 |
| LDA | 86.99 | 82.94 | 83.61 | 84.97 | 69.99 | 85.26 | 87.50 | 82.87 | 83.62 | **85.18** | 70.44 | 85.52 |
| XGBoost | 90.46 | 82.08 | 83.46 | 86.27 | 72.79 | 86.82 | 90.74 | 79.16 | 81.32 | **84.95** | 70.38 | 85.77 |
| LGBM | 90.75 | 81.21 | 82.84 | 85.98 | 72.29 | 86.62 | 89.12 | 80.55 | 82.08 | **84.83** | 69.93 | 85.46 |
| AdaBoost | 87.86 | 81.21 | 82.38 | 84.53 | 69.22 | 85.03 | 88.19 | 80.09 | 81.58 | **84.14** | 68.51 | 84.76 |
| RF | 92.19 | 80.05 | 82.21 | 86.12 | 72.79 | 86.92 | 92.12 | 74.76 | 78.50 | **83.44** | 67.92 | 84.77 |
| RCV | 91.32 | 81.79 | 83.37 | 86.56 | 73.45 | 87.17 | 90.04 | 75.69 | 78.74 | **82.87** | 66.42 | 84.01 |
| Extra Trees | 91.04 | 80.34 | 82.24 | 85.69 | 71.79 | 86.41 | 91.66 | 74.07 | 77.95 | **82.87** | 66.78 | 84.25 |
| Ridge | 87.57 | 81.50 | 82.56 | 84.53 | 69.20 | 84.99 | 84.25 | 76.38 | 78.11 | **80.32** | 60.83 | 81.06 |

However, most ML models demonstrated limited performance overall, likely due to the heterogeneous nature of the input features. The variability in amino acid composition, physicochemical properties, and sequence length introduced complexity that conventional classifiers struggled to capture effectively. The high-dimensional and learned representations were then used to train a broader set of machine learning and deep learning models to improve allergenicity prediction performance.

### 3.5. Feature Selection

Feature extraction is a critical step in developing machine learning-based classifiers for macromolecules, as it enables the transformation of protein sequences into fixed-length numerical representations suitable for computational analysis. In this study, we explored multiple feature generation techniques, including handcrafted features using the Biopython library, contextual embeddings from pre-trained models such as ESM2, ProtFlash, and ProtGPT-2, and binary vectors via one-hot encoding. The performance of these feature sets was evaluated using three classifiers with varying complexity: QDA, RidgeClassifierCV (RCV), and XGBoost, listed in Supplementary Table 1.

On the validation dataset, Biopython-derived features achieved the highest accuracy with QDA (91.26%), followed by 82.87% and 84.95% using RCV and XGBoost, respectively. ESM2 embeddings yielded strong performance with RCV (91.31%) and XGBoost (90.27%), although QDA performed poorly on this representation. One-hot encoded features, by contrast, consistently underperformed, with all classifiers achieving accuracies below 63%. Given their superior performance, we selected Biopython features alongside ESM2-derived embeddings for the downstream model development.

### 3.6. Feature Optimization

We employed dimensionality reduction and feature selection techniques to improve model performance and mitigate issues arising from high-dimensional and potentially redundant features. We used t-distributed stochastic neighbor embedding (t-SNE) to visualize the feature distribution in a two-dimensional space [37,54].

Figure 3a illustrates the original Biopython feature space, where allergenic and non-allergenic sequences exhibit considerable overlap. To enhance class separability, we applied feature selection using a GradientBoosting classifier, which identified the most informative features based on their contribution to classification. The refined Biopython feature set, reduced to 53 dimensions, exhibited improved separation in t-SNE visualization (Figure 3b).

**Figure 3:**
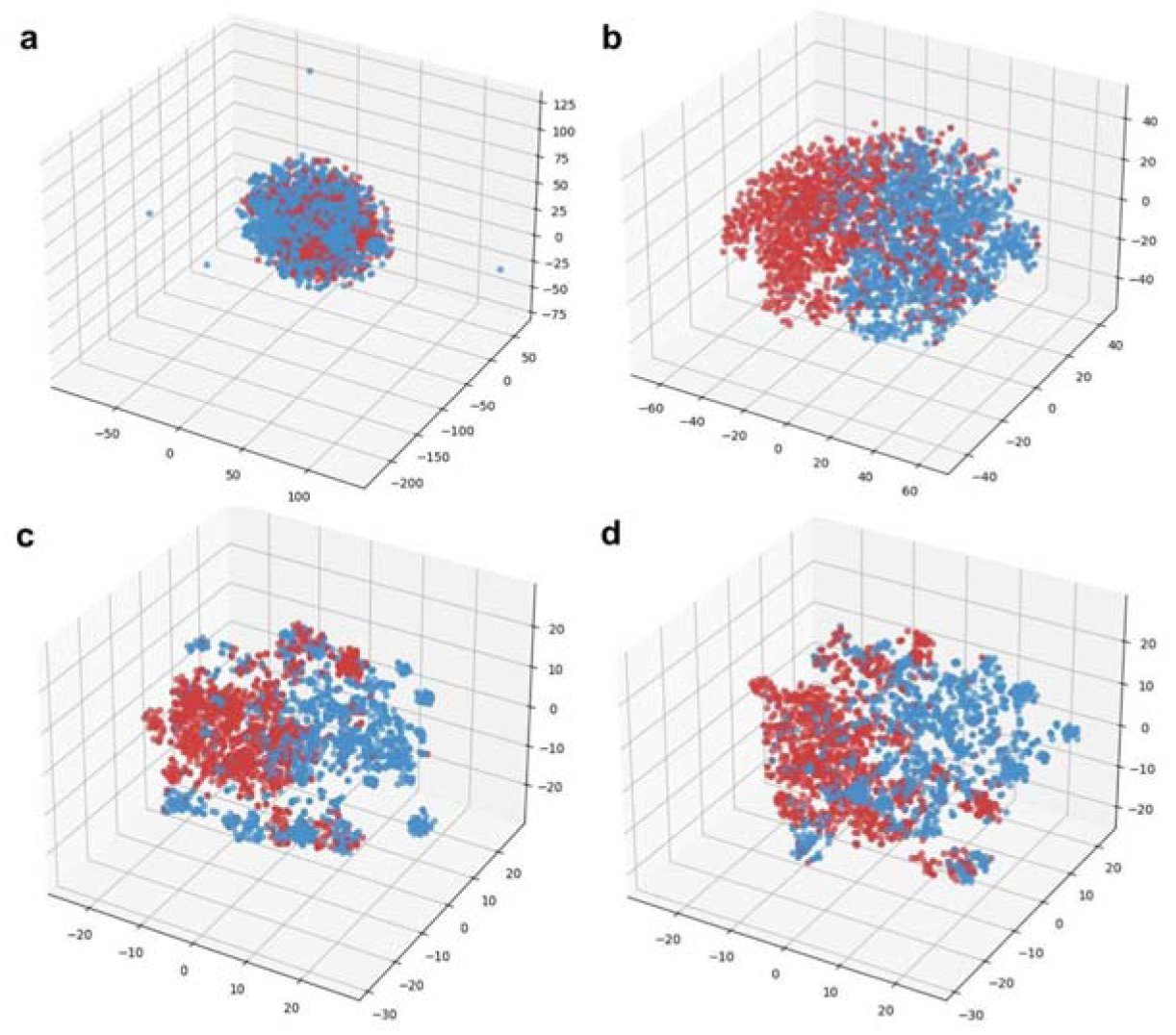
t-SNE visualization of Biopython and ESM2 features before and after feature selection. (a) Raw Biopython feature space (437 features); (b) Biopython features after selection (53 features); (c) Raw ESM2 embeddings (1,280 features); (d) ESM2 features after selection (133 features) using GradientBoosting-based feature importance.

Similarly, we evaluated the 1,280-dimensional ESM2 embeddings (Figure 3c). Although these embeddings provided inherent separation between classes, their high dimensionality posed a risk of overfitting and increased computational complexity. GradientBoosting-based feature selection reduced the feature space to 133 dimensions, enhancing both interpretability and computational efficiency (Figure 3d).

### 3.7. Performance of ESM2 Features on Machine Learning and Deep Learning Models

To evaluate the effectiveness of ESM2-derived embeddings, we implemented five machine learning (ML) models, including: Extremely Randomized Trees (ET), Support Vector Classifier (SVC), k-Nearest Neighbors (KNN), RidgeClassifierCV (RCV), and Light Gradient Boosting Machine (LGBM), alongside three deep learning (DL) architectures: Artificial Neural Network (ANN), Recurrent Neural Network (RNN), and Long Short-Term Memory (LSTM).

Each model was fine-tuned for optimal performance. Among ML models, ET achieved the highest accuracy of 90.28% and an F1 score of 90.45% on the validation set. Among DL models, ANN performed best, achieving 91.71% accuracy and an F1 score of 91.42%. These results indicate that deep contextual embeddings from ESM2 can be effectively leveraged by both traditional and deep learning models for allergenicity classification, as depicted in Table 2.

**Table 2:**
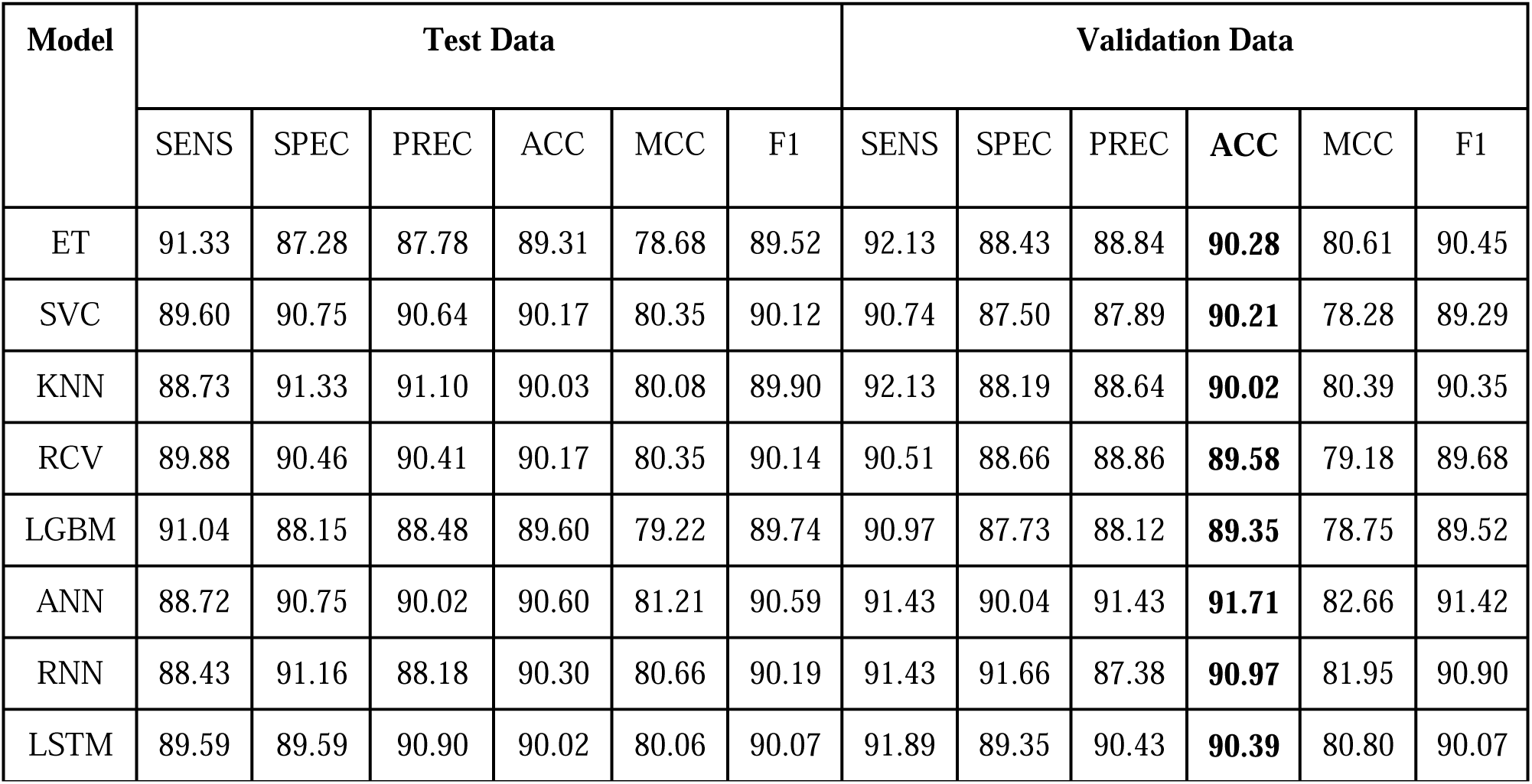
Evaluation of machine learning and deep learning models on ESM2-derived features. Performance is reported on both test and validation datasets using standard classification metrics.

| Model | Test Data |  |  |  |  |  | Validation Data |  |  |  |  |  |
| --- | --- | --- | --- | --- | --- | --- | --- | --- | --- | --- | --- | --- |
|  | SENS | SPEC | PREC | ACC | MCC | F1 | SENS | SPEC | PREC | ACC | MCC | F1 |
| ET | 91.33 | 87.28 | 87.78 | 89.31 | 78.68 | 89.52 | 92.13 | 88.43 | 88.84 | **90.28** | 80.61 | 90.45 |
| SVC | 89.60 | 90.75 | 90.64 | 90.17 | 80.35 | 90.12 | 90.74 | 87.50 | 87.89 | **90.21** | 78.28 | 89.29 |
| KNN | 88.73 | 91.33 | 91.10 | 90.03 | 80.08 | 89.90 | 92.13 | 88.19 | 88.64 | **90.02** | 80.39 | 90.35 |
| RCV | 89.88 | 90.46 | 90.41 | 90.17 | 80.35 | 90.14 | 90.51 | 88.66 | 88.86 | **89.58** | 79.18 | 89.68 |
| LGBM | 91.04 | 88.15 | 88.48 | 89.60 | 79.22 | 89.74 | 90.97 | 87.73 | 88.12 | **89.35** | 78.75 | 89.52 |
| ANN | 88.72 | 90.75 | 90.02 | 90.60 | 81.21 | 90.59 | 91.43 | 90.04 | 91.43 | **91.71** | 82.66 | 91.42 |
| RNN | 88.43 | 91.16 | 88.18 | 90.30 | 80.66 | 90.19 | 91.43 | 91.66 | 87.38 | **90.97** | 81.95 | 90.90 |
| LSTM | 89.59 | 89.59 | 90.90 | 90.02 | 80.06 | 90.07 | 91.89 | 89.35 | 90.43 | **90.39** | 80.80 | 90.07 |

### 3.8. Performance of the Stacked Ensemble Framework

We developed a stacked ensemble model, AllerStack, to further enhance predictive performance, which integrates diverse classifiers trained on heterogeneous features (Biopython-derived and ESM2 embeddings). Stacking combines multiple first-stage base classifiers, whose outputs are input features to a second-stage meta-classifier [36,38,55].

The base classifiers selected for stacking included QDA (91.26% accuracy, F1: 88.23%), ANN (91.71%, F1: 91.42%), and the top-performing ML models—ET (90.28%), SVC (90.21%), and KNN (90.02%). We evaluated seven stacked model configurations (SM1-SM7), combining different subsets of these classifiers. SM6, which included QDA, ANN, SVC, and KNN, achieved the highest performance with an accuracy of 96.87% and an F1 score of 96.86% on validation data, as shown in Table 3.

**Table 3:** Performance of different stacked model configurations using combinations of base classifiers. The seven possible stacked models (SM) were: SM1: QDA, ANN, ET; SM2: QDA, ANN, SVC; SM3: QDA, ANN, KNN; SM4: QDA, ANN, ET, SVC; SM5: QDA, ANN, ET, KNN; SM6: QDA, ANN, SVC, KNN; and SM7: QDA, ANN, ET, SVC, KNN.

| SM | Test Data |  |  |  |  |  | Validation Data |  |  |  |  |  |
| --- | --- | --- | --- | --- | --- | --- | --- | --- | --- | --- | --- | --- |
|  | SENS | SPEC | PREC | ACC | MCC | F1 | SENS | SPEC | PREC | ACC | MCC | F1 |
| **SM6** | **95.66** | **95.37** | **95.38** | **95.52** | **91.04** | **95.52** | **96.52** | **97.22** | **97.20** | **96.87** | **93.75** | **96.86** |
| SM3 | 95.37 | 94.50 | 94.55 | 94.94 | 89.88 | 94.96 | 95.60 | 96.52 | 96.49 | **95.91** | 92.13 | 96.04 |
| SM2 | 96.24 | 95.95 | 95.96 | 96.09 | 92.19 | 96.10 | 95.13 | 96.29 | 96.25 | **95.71** | 91.44 | 95.69 |
| SM4 | 91.32 | 94.50 | 94.32 | 92.91 | 85.88 | 92.80 | 92.36 | 96.52 | 96.37 | **94.44** | 88.96 | 94.32 |
| SM5 | 91.90 | 94.50 | 94.36 | 93.20 | 86.44 | 93.11 | 92.12 | 96.29 | 96.13 | **94.21** | 88.50 | 94.08 |
| SM7 | 91.90 | 94.79 | 94.64 | 93.35 | 86.74 | 93.25 | 92.36 | 95.83 | 95.68 | **94.09** | 88.24 | 93.99 |
| SM1 | 90.75 | 95.08 | 94.86 | 92.91 | 85.91 | 92.76 | 91.89 | 96.06 | 95.89 | **93.98** | 88.03 | 93.85 |

We compared three options for the meta-classifier: logistic regression (LR), XGBoost, and CatBoost. While LR is widely used due to its interpretability [56–59], XGBoost achieved superior performance, likely due to its ability to model non-linear relationships and manage high-dimensional data efficiently. Supplementary Table 2 shows that XGBoost yielded the best results, with a validation accuracy of 96.87% and an F1 score of 96.86%.

Figure 4 illustrates the ROC curves of the base models and the stacked ensemble model. The SM6 configuration achieved the highest area under the ROC curve (AUC = 0.99), indicating superior discrimination between allergenic and non-allergenic proteins. QDA showed the strongest individual performance among base models, followed by SVC and KNN. AllerStack thus demonstrated robust classification capability with high sensitivity (True Positive Rate =96.52%), outperforming all constituent models.

**Figure 4:**
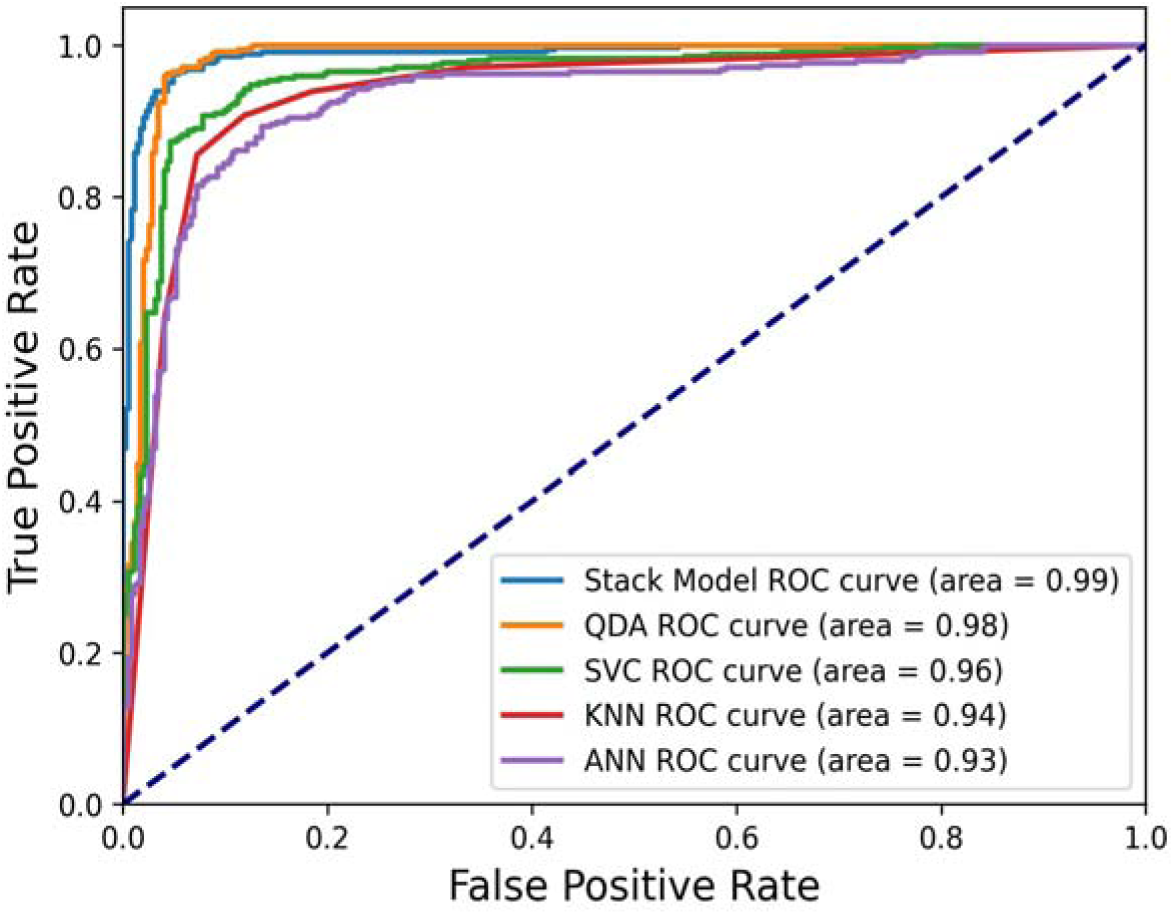
Receiver Operating Characteristic (ROC) curves for base classifiers and the stacked model (SM6). The stacked model exhibits the highest AUC (0.99), indicating superior classification performance compared to individual models.

### 3.9. SHAP Analysis of the Stacked Ensemble Model

SHAP (SHapley Additive exPlanations) is a model-agnostic interpretability method that quantifies the contribution of each feature to individual predictions, enabling transparency in complex machine learning models [60–62]. We applied SHAP analysis to interpret the meta-classifier (XGBoost) and its constituent base models (QDA, SVC, KNN, and ANN) within the AllerStack framework (Figure 5).

**Figure 5:**
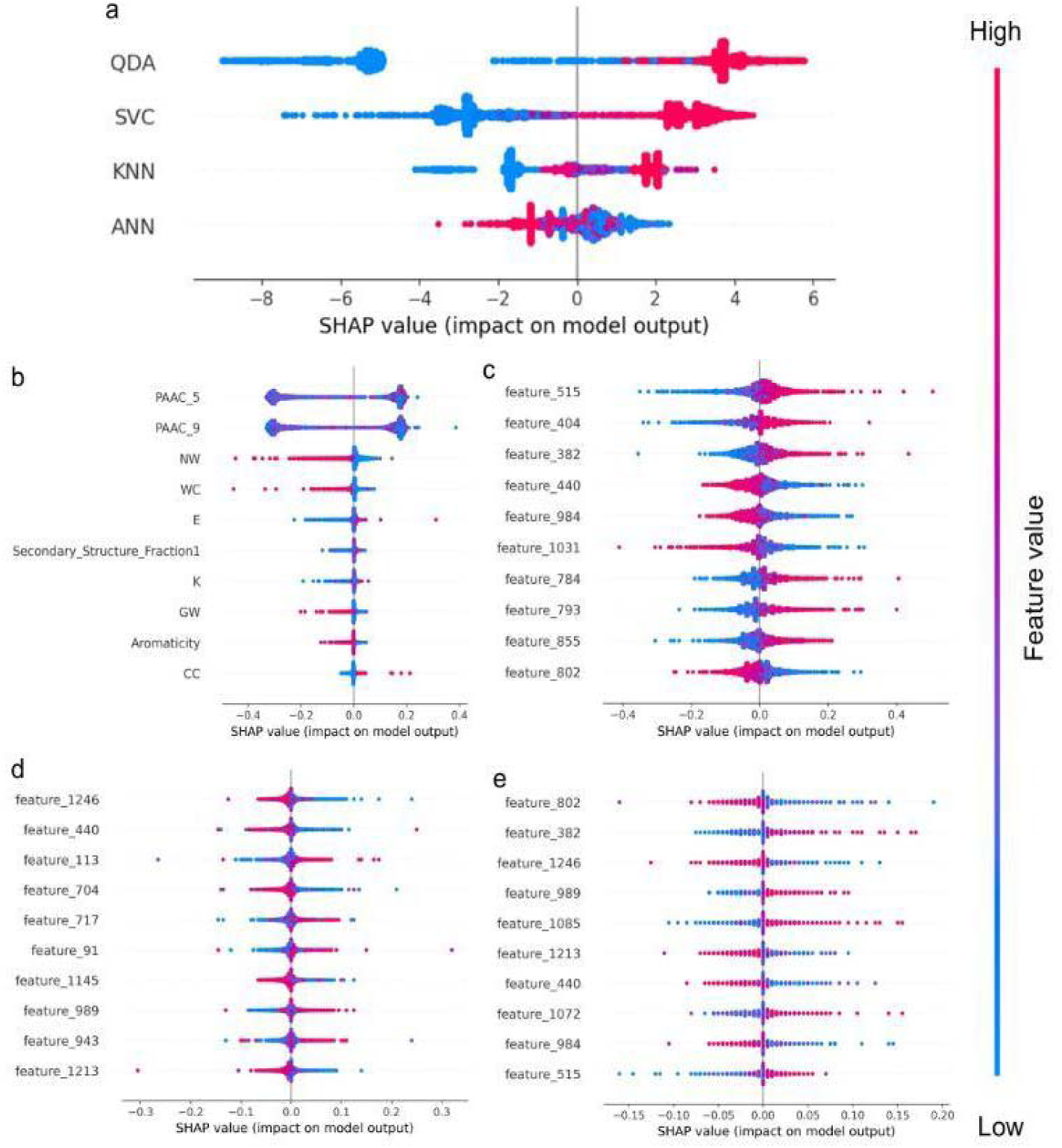
SHAP-based model interpretation of the base classifiers and meta model. (a) Contribution of each base model to the final prediction of XGBoost (meta-model); (b-e) Top 10 contributing features for each base model: (b) QDA (base-model); (c) ANN (base-model); (d) KNN (base-model); (e) SVC (base-model).

Figure 5a highlights the relative contributions of base models to the final prediction. QDA and SVC display a broader range of SHAP values, indicating a more substantial and variable influence on the stacked model’s output compared to KNN and ANN, which was also supported by Figure 4.

Figure 5b-5e depicts the top ten features influencing each base model. For QDA (Figure 5b), key features include PAAC_5, PAAC_9, aromaticity, alpha-helical content (Secondary_Structure_fraction_1), negatively charged residue E, positively charged residue K, and specific dipeptides such as NW, WC, CC, and GW. Notably, feature_440 appears across all three ML base models (SVC, KNN, ANN), while 515, 382, and 802 are shared between SVC and ANN. KNN and ANN commonly use features 989 and 1213. The limited overlap among top features suggests that each classifier captures distinct aspects of the sequence space, reinforcing the value of stacking diverse models to improve generalization.

### 3.10. Performance Comparison with Literature

To contextualize the performance of AllerStack, we compared it against several state-of-the-art allergenicity prediction models. It integrates classical handcrafted features like AAC, DPC, physicochemical descriptors, and deep contextual embeddings from the pre-trained transformer-based ESM2 model.

Unlike most prior models, which are trained on datasets ranging from 4,018 to 20,150 sequences [5,6]. AllerStack was trained and validated on a significantly larger, balanced dataset of 23,860 sequences. This enhanced dataset’s diversity and size contributed to its robust performance.

The model architecture also uses a two-stage stacked ensemble, where predictions from optimized base classifiers serve as meta-classifiers’ features. As shown in Table 4, AllerStack achieved the highest accuracy (96.87%), sensitivity (96.52%), and Matthews Correlation Coefficient (93.75%) among all models. These results reflect its ability to detect allergenic proteins while minimizing false positives and negatives. Additionally, the F1 score of 96.86% underscores the model’s balance between precision and recall, especially in imbalanced datasets. Overall, AllerStack demonstrates improved predictive power, scalability, and interpretability compared to existing allergenicity prediction methods.

**Table 4:** Performance comparison of AllerStack with existing allergenicity prediction. Metrics are based on published literature or reported values.

| Methods | Dataset | SENS | SPEC | PREC | ACC | MCC | F1 | Web server working | Year |
| --- | --- | --- | --- | --- | --- | --- | --- | --- | --- |
| **AllerStack** | **23,860** | **96.52** | **97.22** | **97.20** | **96.87** | **93.75** | **96.86** | **Yes** | **2025** |
| AlgPred 2.0 | 20,150 | 93.10 | 95.36 | NA | 94.23 | 88.04 | NA | Yes | 2021 |
| ProAll-D | 4,854 | NA | NA | 91 | 91.5 | NA | 91 | Yes | 2022 |
| AllerCatPro 2.0 | 4,018 | 93.2 | 98.8 | NA | 96.0 | 92.1 | NA | Yes | 2022 |
| AllerTOPv2 | 4,854 | 86.70 | 90.70 | NA | 88.70 | 77.5 | NA | Yes | 2014 |
| AllerHunter | 14,805 | 83.70 | 96.40 | NA | 95.30 | 73.8 | NA | No | 2009 |
| AllergenFP | 4,854 | 86.80 | 89.10 | NA | 87.90 | 75.9 | NA | Yes | 2014 |

### 3.11. Web Server Interface

We have created a user-friendly web server, AllerStack (https://cosylab.iiitd.edu.in/allerstack/), based on the best stacked model, to predict allergenic proteins. The server enables the user to predict the allergenicity of proteins by providing a single protein sequence or by uploading a TXT, CSV, or FASTA format file as input. This server implements a stacked ensemble approach using AAC, DPC, physicochemical, and ESM2-derived advanced features.

The web server uses a technology stack optimized for efficiency and scalability. The frontend was built with ReactJS, a JavaScript toolkit that allows for the building of dynamic and interactive user interfaces. HTML and CSS were used to structure and style web pages. Flask API endpoints, a lightweight Python web framework, were utilized on the backend. The database is stored in MongoDB, a NoSQL database that enables flexible and scalable data management.

## 4. Conclusions

The accurate prediction of allergenic proteins is critical for preventing allergen exposure through food, pharmaceuticals, and consumer products. With allergies affecting approximately 10–30% of the global population, particularly in developed countries, there is an urgent need for robust computational tools to identify potential allergens before they reach the public domain [42,63,64]. Such predictive models are essential for regulatory compliance and developing hypoallergenic formulations in food and the therapeutic industries [45].

AllerStack is an ensemble-based allergenicity prediction framework trained on a comprehensive dataset of 23,860 allergenic and non-allergenic proteins. By integrating handcrafted features from Biopython (e.g., AAC, DPC, physicochemical properties) with contextual embeddings from the pre-trained ESM2 model, AllerStack effectively captures sequence-level, evolutionary, and structural information. Unlike conventional ML or DL models, it employs a stacked ensemble strategy that leverages the strengths of multiple base learners. This approach achieved a high discriminative performance with an AUC of 0.99, highlighting its reliability in distinguishing allergenic from non-allergenic proteins.

Our findings contribute a scalable and interpretable framework for allergenicity prediction that can support applications in food safety, drug development, and cosmetic product design [65,66]. Future work will explore quantifying allergenic potential, integrating host-specific allergy variability, and expanding to multi-label or multi-class allergen types. AllerStack lays the groundwork for the next generation of predictive models in allergen science and bioinformatics [65–68].

## Acknowledgments

GB thanks Infosys Centre for Artificial Intelligence, Center of Excellence in Healthcare, and Department of Computational Biology, Indraprastha Institute of Information Technology Delhi (IIIT-Delhi) for the computational support. HS is a research scholar in the Complex Systems Laboratory and is thankful to the Department of Science and Technology (DST), Government of India, DST-INSPIRE, for providing the research fellowship.

## CRediT Authorship Contribution Statement

**Harshika Sharma:** Writing – original draft, Writing – review and editing, Project administration, Visualization, Methodology, Formal analysis. **Mansi Goel:** Writing – review and editing, Supervision, Software. **Devkul Sahu:** Data curation. **Mazhar Ayaz Sayed:** Data curation. **Pratham Mangla:** Software. **Pratham Shekhawat:** Software. **Shubham Yadav:** Software. **Ganesh Bagler:** Conceptualization, Investigation, Supervision, Project administration, Writing – review and editing.

## Data and Code Availability

All the datasets and code are available on <u>GitHub</u>.

## Additional Information

See Supplementary information.

## References

[1] B. Singh, A. Karnwal, A. Tripathi, A.K. Upadhyay, Food allergens and related computational biology approaches: a requisite for a healthy life, Bioinforma. Agric. High-Throughput Approaches (2021) 145–160.

[2] A. Navines-Ferrer, E. Serrano-Candelas, G.-J. Molina-Molina, M. Martin, IgE-related chronic diseases and anti-IgE-based treatments, J. Immunol. Res. 2016 (2016) 8163803.

[3] M. Masoli, D. Fabian, S. Holt, R. Beasley, G.I. for Asthma (GINA) Program, The global burden of asthma: executive summary of the GINA Dissemination Committee report, Allergy 59 (2004) 469–478.

[4] B. Anibarro, F.J. Seoane, M. V Mugica, Involvement of hidden allergens in food allergic reactions, J. Investig. Allergol. Clin. Immunol. 17 (2007) 168.

[5] N. Sharma, S. Patiyal, A. Dhall, A. Pande, C. Arora, G.P.S. Raghava, AlgPred 2.0: An improved method for predicting allergenic proteins and mapping of IgE epitopes, Brief. Bioinform. 22 (2021). 10.1093/bib/bbaa294.

[6] H.C. Muh, J.C. Tong, M.T. Tammi, AllerHunter: a SVM-pairwise system for assessment of allergenicity and allergic cross-reactivity in proteins, PLoS One 4 (2009) e5861.

[7] S. Maurer-Stroh, N.L. Krutz, P.S. Kern, V. Gunalan, M.N. Nguyen, V. Limviphuvadh, F. Eisenhaber, G.F. Gerberick, AllerCatPro—prediction of protein allergenicity potential from the protein sequence, Bioinformatics 35 (2019) 3020–3027.

[8] I. Dimitrov, D.R. Flower, I. Doytchinova, AllerTOP-a server for in silico prediction of allergens, in: BMC Bioinformatics, 2013: pp. 1–9.

[9] I. Dimitrov, L. Naneva, I. Doytchinova, I. Bangov, AllergenFP: allergenicity prediction by descriptor fingerprints, Bioinformatics 30 (2014) 846–851.

[10] P.M. Shanthappa, R. Kumar, ProAll-D: protein allergen detection using long short term memory - a deep learning approach, ADMET DMPK 10 (2022) 231–240. 10.5599/admet.1335.

[11] L. Wang, D. Niu, X. Zhao, X. Wang, M. Hao, H. Che, A comparative analysis of novel deep learning and ensemble learning models to predict the allergenicity of food proteins, Foods 10 (2021) 809.

[12] C. He, X. Ye, Y. Yang, L. Hu, Y. Si, X. Zhao, L. Chen, Q. Fang, Y. Wei, F. Wu, others, DeepAlgPro: an interpretable deep neural network model for predicting allergenic proteins, Brief. Bioinform. 24 (2023) bbad246.

[13] F.M. Garcia-Moreno, M.A. Gutiérrez-Naranjo, ALLERDET: a novel web app for prediction of protein allergenicity, J. Biomed. Inform. 135 (2022) 104217.

[14] M.R. Islam, M.A. Kashem, L. Mia, AllerHybrid: A Hybrid System to Predict the Allergen Using K-mer and Physicochemical Properties, in: 2021 5th Int. Conf. Electr. Eng. Inf. Commun. Technol., 2021: pp. 1–6.

[15] A. Kumar, P.S. Rana, A deep learning based ensemble approach for protein allergen classification, PeerJ Comput. Sci. 9 (2023) e1622.

[16] R. van Ree, D. Sapiter Ballerda, M.C. Berin, L. Beuf, A. Chang, G. Gadermaier, P.A. Guevera, K. Hoffmann-Sommergruber, E. Islamovic, L. Koski, J. Kough, G.S. Ladics, S. McClain, K.A. McKillop, S. Mitchell-Ryan, C.A. Narrod, L. Pereira Mouriès, S. Pettit, L.K. Poulsen, A. Silvanovich, P. Song, S.S. Teuber, C. Bowman, The COMPARE Database: A Public Resource for Allergen Identification, Adapted for Continuous Improvement, Front. Allergy 2 (2021) 700533. 10.3389/FALGY.2021.700533/BIBTEX.

[17] R.E. Goodman, M. Ebisawa, F. Ferreira, H.A. Sampson, R. van Ree, S. Vieths, J.L. Baumert, B. Bohle, S. Lalithambika, J. Wise, S.L. Taylor, AllergenOnline: A peer-reviewed, curated allergen database to assess novel food proteins for potential cross-reactivity, Mol. Nutr. Food Res. 60 (2016) 1183–1198. 10.1002/mnfr.201500769.

[18] A. Mari, C. Rasi, P. Palazzo, E. Scala, Allergen databases: Current status and perspectives, Curr. Allergy Asthma Rep. 9 (2009) 376–383. 10.1007/s11882-009-0055-9.

[19] S. Sudharson, T. Kalic, C. Hafner, H. Breiteneder, Newly defined allergens in the WHO/IUIS Allergen Nomenclature Database during 01/2019-03/2021, Allergy Eur. J. Allergy Clin. Immunol. 76 (2021) 3359–3373. 10.1111/all.15021.

[20] M.W.E.J. Fiers, G.A. Kleter, H. Nijland, A.A.C.M. Peijnenburg, J.P. Nap, R.C.H.J. van Ham, Allermatch^TM^, a webtool for the prediction of potential allergenicity according to current FAO/WHO Codex alimentarius guidelines, BMC Bioinformatics 5 (2004) 1–6. 10.1186/1471-2105-5-133/TABLES/2.

[21] K. Kadam, R. Karbhal, V.K. Jayaraman, S. Sawant, U. Kulkarni-Kale, AllerBase: a comprehensive allergen knowledgebase, Database 2017 (2017) 1–12. 10.1093/DATABASE/BAX066.

[22] C.H. Schein, Identifying Similar Allergens and Potentially Cross-Reacting Areas Using Structural Database of Allergenic Proteins (SDAP) Tools and D-Graph, Methods Mol. Biol. 2717 (2024) 269–284. 10.1007/978-1-0716-3453-0_18.

[23] allergen in UniProtKB search (1286) | UniProt, (n.d.). https://www.uniprot.org/uniprotkb?query=allergen&facets=reviewed%3Atrue (accessed August 23, 2024).

[24] NOT allergen in UniProtKB search (570578) | UniProt, (n.d.). https://www.uniprot.org/uniprotkb?query=NOT+allergen&facets=reviewed%3Atrue (accessed August 23, 2024).

[25] M. De Angelis, G. Garruti, F. Minervini, L. Bonfrate, P. Portincasa, M. Gobbetti, The Food-gut Human Axis: The Effects of Diet on Gut Microbiota and Metabolome, Curr. Med. Chem. 26 (2017) 3567–3583. 10.2174/0929867324666170428103848.

[26] L. Fu, B. Niu, Z. Zhu, S. Wu, W. Li, CD-HIT: accelerated for clustering the next-generation sequencing data, Bioinformatics 28 (2012) 3150–3152. 10.1093/BIOINFORMATICS/BTS565.

[27] P.J.A. Cock, T. Antao, J.T. Chang, B.A. Chapman, C.J. Cox, A. Dalke, I. Friedberg, T. Hamelryck, F. Kauff, B. Wilczynski, M.J.L. De Hoon, Biopython: freely available Python tools for computational molecular biology and bioinformatics, Bioinformatics 25 (2009) 1422. 10.1093/BIOINFORMATICS/BTP163.

[28] P.K. Meher, T.K. Sahu, V. Saini, A.R. Rao, Predicting antimicrobial peptides with improved accuracy by incorporating the compositional, physico-chemical and structural features into Chou’s general PseAAC, Sci. Reports 2017 71 7 (2017) 1–12. 10.1038/srep42362.

[29] J.V. Kringelum, M. Nielsen, S.B. Padkjær, O. Lund, Structural analysis of B-cell epitopes in antibody:protein complexes, Mol. Immunol. 53 (2013) 24–34. 10.1016/J.MOLIMM.2012.06.001.

[30] E. Öztemiz Topcu, G. Gadermaier, To stay or not to stay intact as an allergen: the endolysosomal degradation assay used as tool to analyze protein immunogenicity and T cell epitopes, Front. Allergy 5 (2024) 1440360. 10.3389/FALGY.2024.1440360/BIBTEX.

[31] K.M. Makwana, R. Mahalakshmi, Implications of aromatic–aromatic interactions: From protein structures to peptide models, Protein Sci. 24 (2015) 1920–1933. 10.1002/PRO.2814.

[32] A. Rives, J. Meier, T. Sercu, S. Goyal, Z. Lin, J. Liu, D. Guo, M. Ott, C.L. Zitnick, J. Ma, R. Fergus, Biological structure and function emerge from scaling unsupervised learning to 250 million protein sequences, Proc. Natl. Acad. Sci. U. S. A. 118 (2021) e2016239118. 10.1073/PNAS.2016239118/SUPPL_FILE/PNAS.2016239118.SAPP.PDF.

[33] Y. Zhang, X. Chen, B. Jin, S. Wang, S. Ji, W. Wang, J. Han, A Comprehensive Survey of Scientific Large Language Models and Their Applications in Scientific Discovery, (2024). https://arxiv.org/abs/2406.10833v1 (accessed August 23, 2024).

[34] D. Upadhyay, J. Manero, M. Zaman, S. Sampalli, Gradient Boosting Feature Selection with Machine Learning Classifiers for Intrusion Detection on Power Grids, IEEE Trans. Netw. Serv. Manag. 18 (2021) 1104–1116. 10.1109/TNSM.2020.3032618.

[35] Z. Xu, G. Huang, K.Q. Weinberger, A.X. Zheng, Gradient boosted feature selection, Proc. ACM SIGKDD Int. Conf. Knowl. Discov. Data Min. (2014) 522–531. 10.1145/2623330.2623635.

[36] A.I. Albu, M.I. Bocicor, G. Czibula, MM-StackEns: A new deep multimodal stacked generalization approach for protein–protein interaction prediction, Comput. Biol. Med. 153 (2023) 106526. 10.1016/J.COMPBIOMED.2022.106526.

[37] C. Lei, Z. Lu, M. Wang, M. Li, StackCPA: A stacking model for compound-protein binding affinity prediction based on pocket multi-scale features, Comput. Biol. Med. 164 (2023) 107131. 10.1016/J.COMPBIOMED.2023.107131.

[38] Y. Chen, L. Gao, T. Zhang, Stack-VTP: prediction of vesicle transport proteins based on stacked ensemble classifier and evolutionary information, BMC Bioinformatics 24 (2023) 1–18. 10.1186/S12859-023-05257-5/TABLES/7.

[39] M.N. Nguyen, N.L. Krutz, V. Limviphuvadh, A.L. Lopata, G.F. Gerberick, S. Maurer-Stroh, AllerCatPro 2.0: a web server for predicting protein allergenicity potential, Nucleic Acids Res. 50 (2022). 10.1093/nar/gkac446.

[40] I. Dimitrov, I. Bangov, D.R. Flower, AllerTOP v.2-a server for in silico prediction of allergens, (n.d.). 10.1007/s00894-014-2278-5.

[41] C. Radauer, H. Breiteneder, Evolutionary biology of plant food allergens, J. Allergy Clin. Immunol. 120 (2007) 518–525. 10.1016/J.JACI.2007.07.024.

[42] S.H. Sicherer, H.A. Sampson, Food allergy: Epidemiology, pathogenesis, diagnosis, and treatment, J. Allergy Clin. Immunol. 133 (2014) 291–307.e5. 10.1016/J.JACI.2013.11.020.

[43] A.M. Chiriac, Y. Wang, R. Schrijvers, P.J. Bousquet, T. Mura, N. Molinari, P. Demoly, Designing Predictive Models for Beta-Lactam Allergy Using the Drug Allergy and Hypersensitivity Database, J. Allergy Clin. Immunol. Pract. 6 (2018) 139–148.e2. 10.1016/j.jaip.2017.04.045.

[44] H. Breiteneder, E.N.C. Mills, Molecular properties of food allergens, J. Allergy Clin. Immunol. 115 (2005) 14–23. 10.1016/J.JACI.2004.10.022.

[45] H. Djeghim, I. Bellil, O. Benslama, S. Lekmine, E. Temim, H. Boufendi, I. Postigo, P. Sánchez, D. Khelifi, Effects of genetic diversity on the allergenicity of peanut (Arachis hypogaea) proteins: identification of the hypoallergenic accessions using BALB/c mice model and in silico analysis of Ara h 3 allergen cross-reactivity, J. Proteomics 306 (2024) 105264. 10.1016/J.JPROT.2024.105264.

[46] R. Sharma, A.K. Singh, V. Umashankar, Characterization of allergenic epitopes of Ory s1 protein from Oryza sativa and its homologs, Bioinformation 4 (2009) 12. 10.6026/97320630004012.

[47] W.J. Pichler, A. Beeler, M. Keller, M. Lerch, S. Posadas, D. Schmid, Z. Spanou, A. Zawodniak, B. Gerber, Pharmacological Interaction of Drugs with Immune Receptors: The p-i Concept, Allergol. Int. 55 (2006) 17–25. 10.2332/ALLERGOLINT.55.17.

[48] Q. Geng, Y. Zhang, M. Song, X. Zhou, Y. Tang, Z. Wu, H. Chen, Allergenicity of peanut allergens and its dependence on the structure, Compr. Rev. Food Sci. Food Saf. 22 (2023) 1058–1081. 10.1111/1541-4337.13101.

[49] J. Cui, L.Y. Han, H. Li, C.Y. Ung, Z.Q. Tang, C.J. Zheng, Z.W. Cao, Y.Z. Chen, Computer prediction of allergen proteins from sequence-derived protein structural and physicochemical properties, Mol. Immunol. 44 (2007) 514–520. 10.1016/J.MOLIMM.2006.02.010.

[50] J. Pekar, D. Ret, E. Untersmayr, Stability of allergens, Mol. Immunol. 100 (2018) 14–20. 10.1016/J.MOLIMM.2018.03.017.

[51] A. Cianferoni, J.M. Spergel, Food Allergy: Review, Classification and Diagnosis, Allergol. Int. 58 (2009) 457–466. 10.2332/ALLERGOLINT.09-RAI-0138.

[52] K.D. Collins, G.W. Neilson, J.E. Enderby, Ions in water: Characterizing the forces that control chemical processes and biological structure, Biophys. Chem. 128 (2007) 95–104. 10.1016/J.BPC.2007.03.009.

[53] T.L. Bailey, STREME: accurate and versatile sequence motif discovery, Bioinformatics 37 (2021) 2834–2840. 10.1093/BIOINFORMATICS/BTAB203.

[54] A. Sharma, A. Lysenko, K.A. Boroevich, E. Vans, T. Tsunoda, DeepFeature: feature selection in nonimage data using convolutional neural network, Brief. Bioinform. 22 (2021) 1–12. 10.1093/BIB/BBAB297.

[55] C. Chen, Q. Zhang, B. Yu, Z. Yu, P.J. Lawrence, Q. Ma, Y. Zhang, Improving protein-protein interactions prediction accuracy using XGBoost feature selection and stacked ensemble classifier, Comput. Biol. Med. 123 (2020) 103899. 10.1016/J.COMPBIOMED.2020.103899.

[56] A. Bashir, M. Mohammed, F. Mussallum, M. Khalafallah Elbashir, A.B. Musa, A. Mussallum, SVM and Naïve Bayes Stacking Approach for Improving Gene Expression Data Classification Using Logistic Regression, Artic. Int. J. Adv. Soft Comput. Its Appl. 13 (2021). https://www.researchgate.net/publication/355960625 (accessed August 23, 2024).

[57] M. Samieinasab, S.A. Torabzadeh, A. Behnam, A. Aghsami, F. Jolai, Meta-Health Stack: A new approach for breast cancer prediction, Healthc. Anal. 2 (2022) 100010. 10.1016/J.HEALTH.2021.100010.

[58] P. Rajendra, S. Latifi, Prediction of diabetes using logistic regression and ensemble techniques, Comput. Methods Programs Biomed. Updat. 1 (2021) 100032. 10.1016/J.CMPBUP.2021.100032.

[59] J. Liu, X. Dong, H. Zhao, Y. Tian, Predictive Classifier for Cardiovascular Disease Based on Stacking Model Fusion, Process. 2022, Vol. 10, Page 749 10 (2022) 749. 10.3390/PR10040749.

[60] H. Chen, S. Lundberg, S.I. Lee, Explaining Models by Propagating Shapley Values of Local Components, Stud. Comput. Intell. 914 (2021) 261–270. 10.1007/978-3-030-53352-6_24.

[61] H. Sahlaoui, E.A.A. Alaoui, A. Nayyar, S. Agoujil, M.M. Jaber, Predicting and Interpreting Student Performance Using Ensemble Models and Shapley Additive Explanations, IEEE Access 9 (2021) 152688–152703. 10.1109/ACCESS.2021.3124270.

[62] Y. Nohara, K. Matsumoto, H. Soejima, N. Nakashima, Explanation of machine learning models using shapley additive explanation and application for real data in hospital, Comput. Methods Programs Biomed. 214 (2022) 106584. 10.1016/J.CMPB.2021.106584.

[63] S. Prescott, K.J. Allen, Food allergy: Riding the second wave of the allergy epidemic, Pediatr. Allergy Immunol. 22 (2011) 155–160. 10.1111/J.1399-3038.2011.01145.X.

[64] W. Loh, M.L.K. Tang, The Epidemiology of Food Allergy in the Global Context, Int. J. Environ. Res. Public Heal. 2018, Vol. 15, Page 2043 15 (2018) 2043. 10.3390/IJERPH15092043.

[65] J.M. Larsen, C.H. Bang-Berthelsen, K. Qvortrup, A.I. Sancho, A.H. Hansen, K.I.H. Andersen, S.S.N. Thacker, T. Eiwegger, J. Upton, K.L. Bøgh, Production of allergen-specific immunotherapeutic agents for the treatment of food allergy, Crit. Rev. Biotechnol. 40 (2020) 881–894. 10.1080/07388551.2020.1772194.

[66] X. Pi, Y. Sun, J. Cheng, G. Fu, M. Guo, A review on polyphenols and their potential application to reduce food allergenicity, Crit. Rev. Food Sci. Nutr. 63 (2023) 10014– 10031. 10.1080/10408398.2022.2078273.

[67] J.S. Kandula, V.P.K. Rayala, R. Pullapanthula, Chirality: An inescapable concept for the pharmaceutical, bio-pharmaceutical, food, and cosmetic industries, Sep. Sci. Plus 6 (2023) 2200131. 10.1002/SSCP.202200131.

[68] S. Gajbhiye, K. Pal, Toxic and Allergic Responses Caused by Secondary Metabolites Used in Cosmetic Formulations, Bioprospecting Nat. Sources Cosmeceuticals (2024) 73–104. 10.1039/9781837672288-00073.

