## Supplementary material for "AllerStack: Predicting Allergenic Proteins with a Stacked Ensemble Approach": Supplemenatry Data

1. **Compositional Analysis**

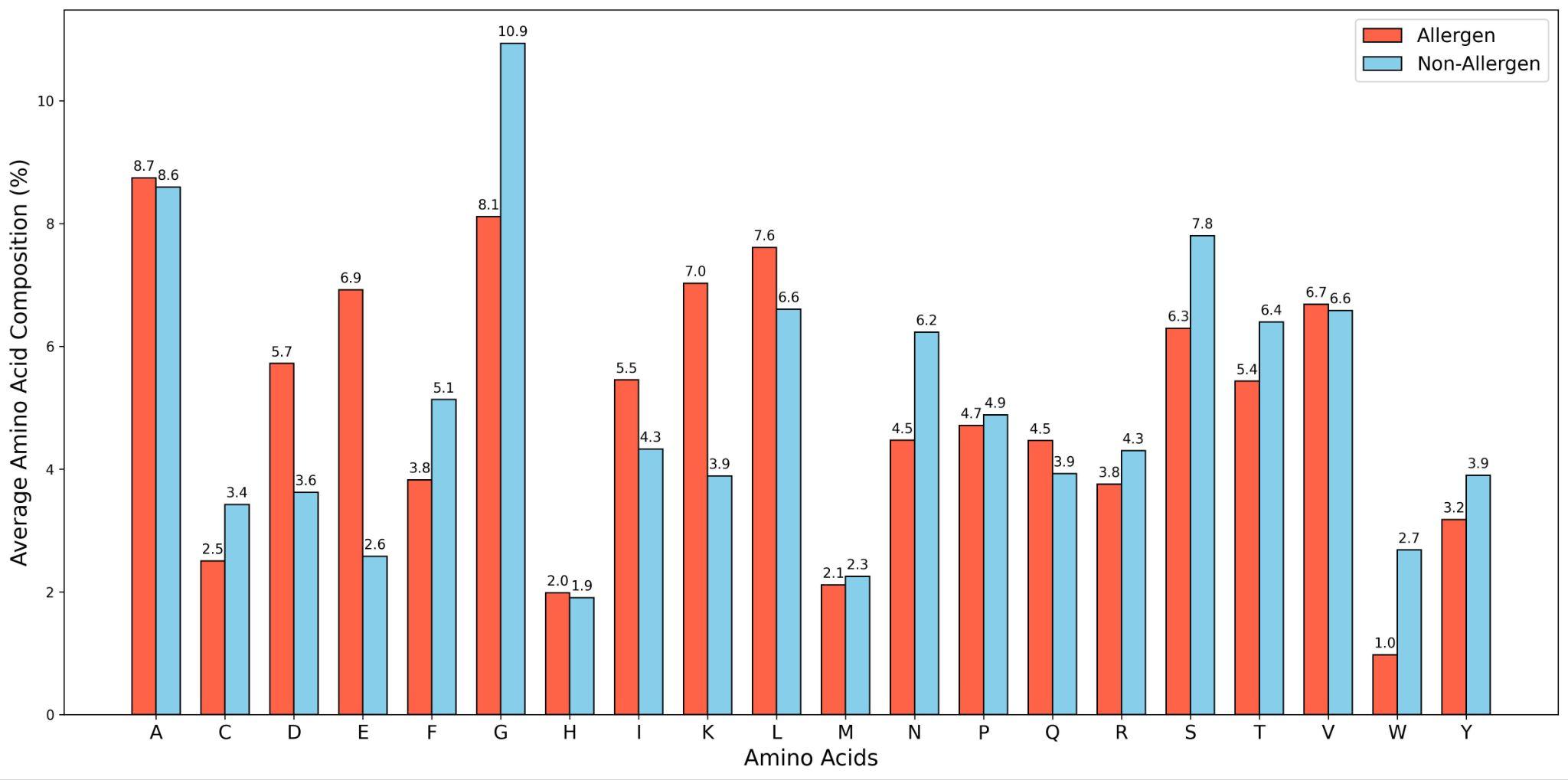

**Supplementary Figure 1:** Average amino acid composition (%) of the allergens and non-allergens. Amino acid alanine (A) has the highest allergen, while glycine (G) has the highest non-allergen. Moreover, it is seen that amino acids like Aspartic Acid (D), Glutamic Acid (E), Isoleucine (I), Lysine (K), Leucine (L), and Glutamine (Q) are more prominent in allergenic sequences, whereas cysteine (C), phenylalanine (F), glycine (G), asparagine (N), arginine (R), serine (S), threonine (T), and tryptophan (W) are more in non-allergenic sequences.

2. **Physicochemical Analysis**

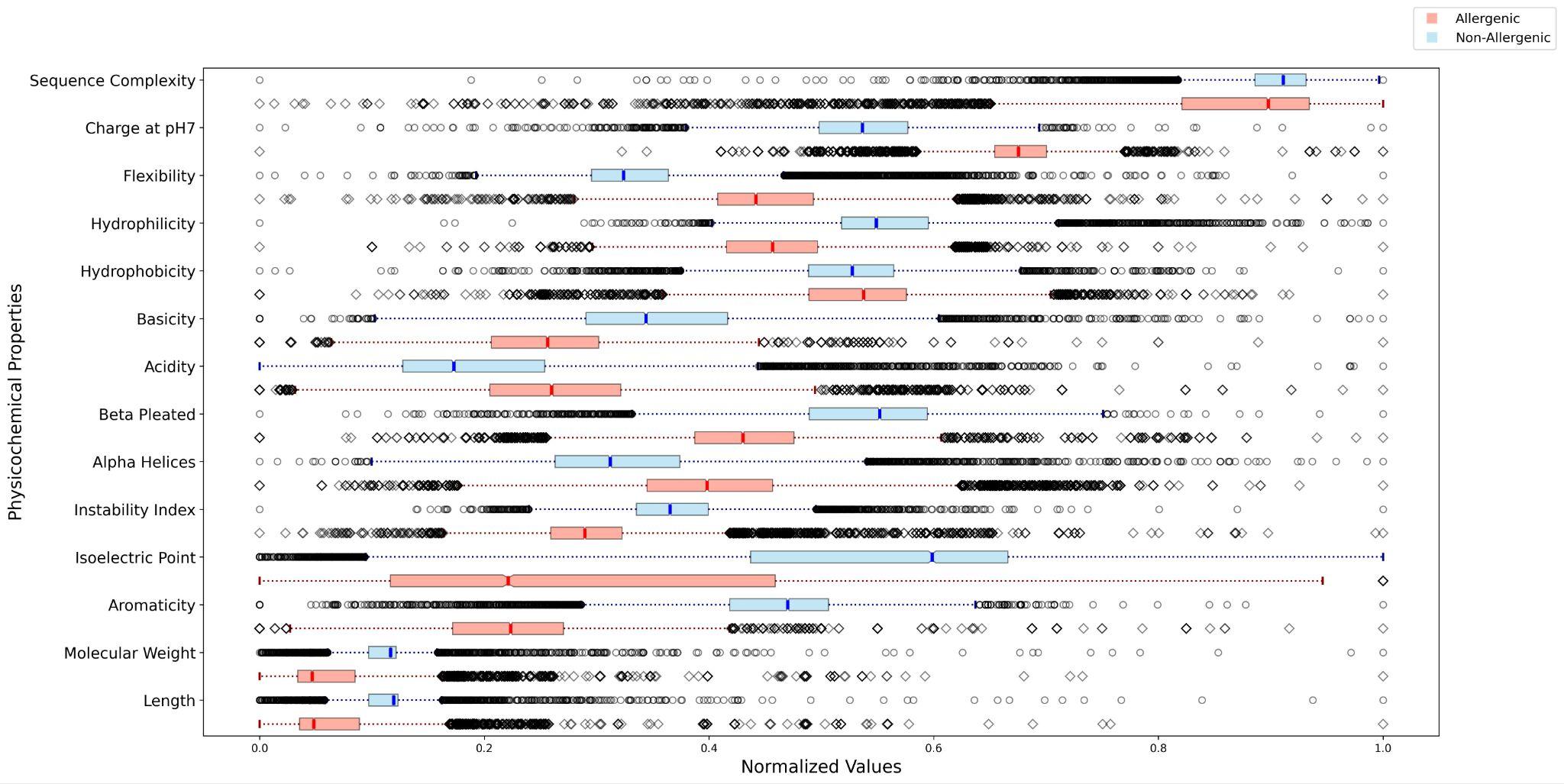

**Supplementary Figure 2:** Physicochemical analysis of the allergenic and non-allergenic proteins. The figure shows the comparison of molecular weight, length, aromaticity, isoelectric point (pI), instability index, alpha helices, beta pleated sheets, acidity, basicity, hydrophobicity, hydrophilicity, flexibility, charge at pH 7, and sequence complexity.

3. **Feature Extraction**

| **Feature** | **Model** | **Test Data** | | | | | | **Validation Data** | | | | | |
| --- | --- | --- | --- | --- | --- | --- | --- | --- | --- | --- | --- | --- | --- |
|  |  | SENS | SPEC | PREC | ACC | MCC | F1 | SENS | SPEC | PREC | **ACC** | MCC | F1 |
| Bio-  python | QDA | 95.66 | 92.07 | 93.22 | 90.83 | 84.75 | 89.78 | 94.52 | 91.23 | 94.23 | **91.26** | 86.58 | 88.23 |
|  | RCV | 91.32 | 81.79 | 83.37 | 86.56 | 73.45 | 87.17 | 90.04 | 75.69 | 78.74 | **82.87** | 66.42 | 84.01 |
|  | XGB | 90.46 | 82.08 | 83.46 | 86.27 | 72.79 | 86.82 | 90.74 | 79.16 | 81.32 | **84.95** | 70.38 | 85.77 |
| ESM2 | QDA | 95.37 | 45.95 | 63.83 | 70.66 | 47.54 | 76.47 | 96.06 | 47.22 | 64.54 | **71.64** | 49.60 | 77.20 |
|  | RCV | 89.88 | 89.59 | 89.62 | 89.74 | 79.48 | 89.75 | 91.43 | 91.20 | 91.22 | **91.31** | 82.63 | 91.32 |
|  | XGB | 91.32 | 88.15 | 88.51 | 89.74 | 79.52 | 89.90 | 91.66 | 88.88 | 89.18 | **90.27** | 80.58 | 90.41 |
| Prot-  flash | QDA | 92.48 | 70.80 | 76.01 | 81.64 | 64.83 | 83.44 | 93.28 | 69.44 | 75.32 | **81.33** | 64.59 | 83.35 |
|  | RCV | 85.83 | 86.12 | 86.08 | 85.98 | 71.96 | 85.96 | 86.80 | 84.72 | 85.03 | **85.76** | 71.54 | 85.91 |
|  | XGB | 84.39 | 85.83 | 85.63 | 85.11 | 70.23 | 85.00 | 87.26 | 84.72 | 85.10 | **85.99** | 72.01 | 86.17 |
| Prot-  gpt2 | QDA | 94.22 | 66.18 | 73.58 | 80.20 | 62.28 | 82.63 | 94.44 | 64.35 | 72.59 | **79.39** | 61.65 | 82.09 |
|  | RCV | 81.21 | 87.57 | 86.72 | 84.39 | 68.92 | 83.88 | 86.80 | 87.50 | 87.41 | **87.15** | 74.43 | 87.10 |
|  | XGB | 89.01 | 86.12 | 86.51 | 87.57 | 75.17 | 87.77 | 90.04 | 85.64 | 86.25 | **87.84** | 75.76 | 88.10 |
| One-  hot | QDA | 30.92 | 91.04 | 77.53 | 60.98 | 27.48 | 44.21 | 35.18 | 84.72 | 69.72 | **59.95** | 22.91 | 46.76 |
|  | RCV | 30.34 | 93.35 | 82.03 | 61.85 | 30.51 | 44.30 | 35.18 | 88.42 | 75.24 | **61.80** | 27.89 | 47.95 |
|  | XGB | 30.63 | 92.77 | 80.91 | 61.70 | 29.87 | 44.44 | 35.41 | 89.12 | 76.50 | **62.26** | 29.08 | 48.41 |

**Supplementary Table 1:** Performance of various features (Biopython, ESM2, ProtFlash, ProtGPT2, and One-hot encoding) using different classifiers (QDA, RCV, XGBoost). As shown in Table ESM2 and Biopython, the features are performing well, with an ACC of greater than 90% in at least one of the three models.

4. **Performance of the Stack-Model Framework**

| **Meta Model** | **Test Data** | | | | | | **Validation Data** | | | | | |
| --- | --- | --- | --- | --- | --- | --- | --- | --- | --- | --- | --- | --- |
|  | SENS | SPEC | PREC | ACC | MCC | F1 | SENS | SPEC | PREC | **ACC** | MCC | F1 |
| XGBoost Classifier | 95.66 | 95.37 | 95.38 | 95.52 | 91.04 | 95.52 | 96.52 | 97.22 | 97.20 | **96.87** | 93.75 | 96.86 |
| CatBoost Classifier | 95.08 | 91.90 | 92.15 | 93.49 | 87.03 | 93.59 | 94.67 | 90.04 | 90.48 | **92.36** | 84.81 | 92.53 |
| LR Classifier | 94.21 | 91.61 | 91.83 | 92.91 | 85.86 | 93.01 | 93.51 | 91.43 | 91.60 | **92.47** | 84.97 | 92.55 |

**Supplementary Table 2:** Performance of various meta-models on the stack model. XGBoost Classifier performs best, with an accuracy of 96.87% and F1 score of 96.86%.
